# Acute hyper- and hypoglycemia uncouples the metabolic cooperation between glucose and lactate to disrupt sleep

**DOI:** 10.1101/2022.09.15.507967

**Authors:** Caitlin M. Carroll, Molly Stanley, Ryan V. Raut, Nicholas J. Constantino, Riley E. Irmen, Anish Mitra, J. Andy Snipes, Marcus E. Raichle, David M. Holtzman, Robert W. Gould, Kenneth T. Kishida, Shannon L. Macauley

**Author notes:** **Corresponding author** Corresponding author: Shannon L. Macauley, PhD, Associate Professor, Department of Physiology & Pharmacology, 575 N. Patterson Ave. Winston-Salem, NC 27101. Co-first authors (Caitlin M. Carroll, Molly Stanley).

## Abstract

The sleep-wake cycle is a master regulator of metabolic and neuronal activity and when altered, can have profound effects on metabolic health and disease. Although consideration is given to how fluctuations in blood glucose affect peripheral physiology and metabolism, less is known about how glucose dysregulation impacts the intrinsic cooperation between brain metabolism and neuronal activity to regulate sleep. To understand the effect of peripheral hyper- and hypoglycemia on these relationships, we paired biosensors measuring hippocampal interstitial fluid (ISF) levels of glucose and lactate with cortical EEG/EMG recordings to produce simultaneous subsecond recordings of ISF glucose, lactate, and sleep-wake states. First, we describe a conserved temporal relationships between ISF glucose and lactate based on their intrinsic oscillations, diurnal rhythms, and sleep/wake cycles. ISF glucose and lactate oscillations are largely anti-correlated but the frequency of their oscillations dictate their power, coherence, and phase. While ISF glucose and lactate both have diurnal fluctuations, only ISF lactate is consistently elevated during wake. During wake, fluctuations in ISF lactate are associated with changes in the EEG power spectrum, suggesting wake-related activity is more closely associated with ISF lactate. Modulation of glucose availability via both hyper- or hypoglycemia disrupts the relationship between peripheral metabolism, brain metabolism, and sleep. Hyper- and hypo-glycemia increase ISF lactate, decrease NREM, and alter EEG spectral activity, again demonstrating ISF lactate drives wake-associated behaviors and disrupts sleep. Taken together, these studies demonstrate that peripheral glucose homeostasis is necessary for maintaining the relationships between brain metabolism, neuronal activity, and sleep-wake patterns and deviations in blood glucose levels are sufficient to disrupt the metabolic signature of sleep-wake states, putting the brain at risk in diseases like type-2-diabetes and Alzheimer’s disease.

**Graphical Abstract. Peripheral glucose homeostasis directly modifies sleep/wake patterns:** 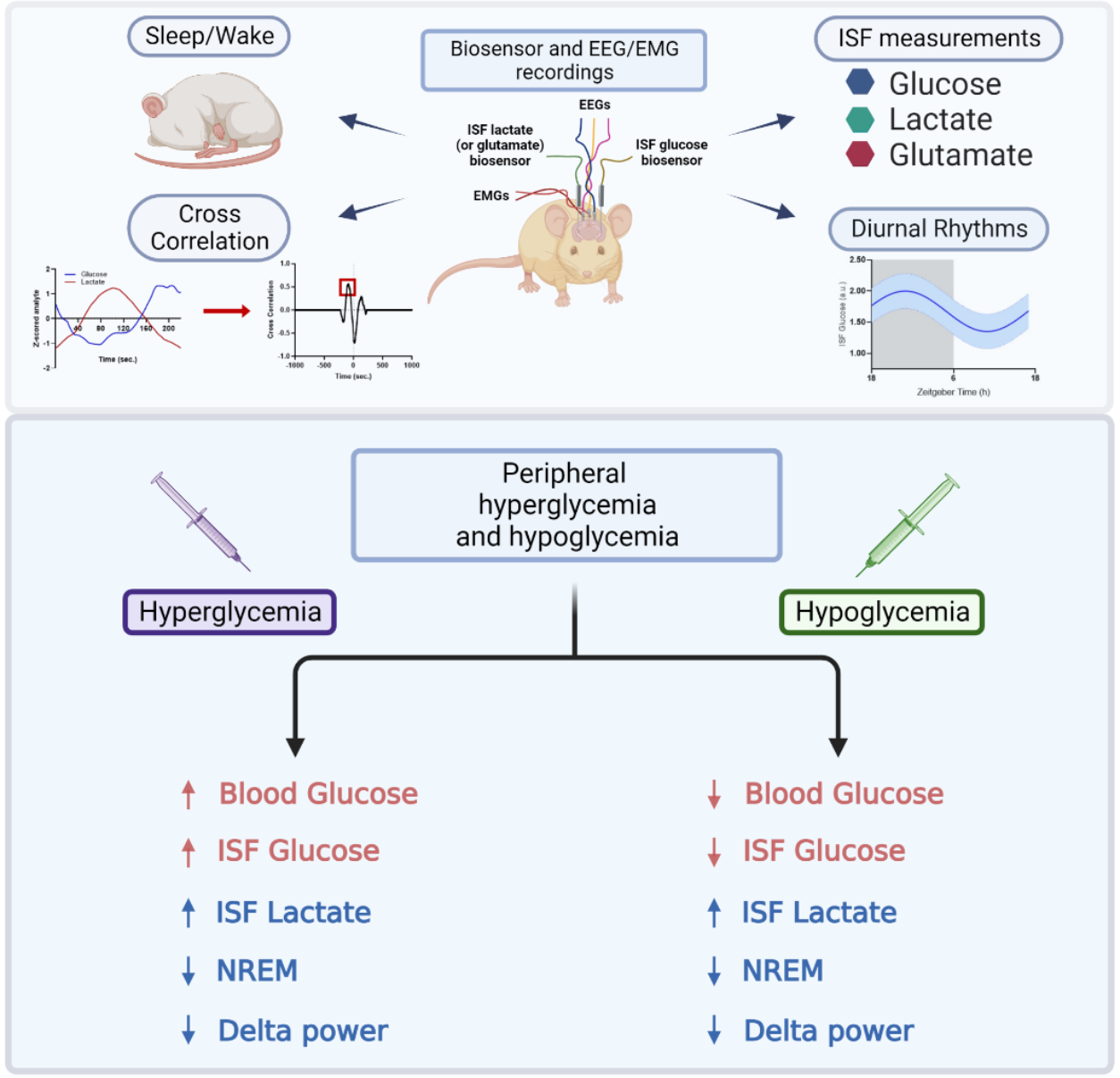

## Introduction

Sleep and metabolic health are profoundly interconnected, where alterations in sleep fitness negatively impact metabolic function. The prevalence of sleep disturbances continues to grow steadily, with nearly 1/3 of US adults currently reporting fewer than 6 hours of sleep a night (1, 2). While poor sleep causes many adverse health outcomes (6), the association with metabolic disease is particularly important given the concurrent, rapid rise in rates of type-2-diabetes (T2D) (3). Disruptions to sleep quality and quantity are associated with increased risk of T2D (2, 4, 5). Suppression of slow wave sleep, the restorative component of non-rapid eye movement (NREM), is associated with glucose intolerance (6-8), insulin resistance (9), and T2D (10). Moreover, individuals with T2D have worse sleep efficiency and greater sleep fragmentation compared to metabolically healthy individuals (11). Taken together, a clear relationship exists between sleep and metabolic homeostasis, but the mechanisms connecting peripheral metabolic changes to altered sleep-wake patterns remains poorly characterized.

In the healthy brain, interstitial fluid (ISF) levels of glucose and lactate are intrinsically coupled and reflect the dynamic relationship between cerebral metabolism and neuronal activity. While the brain represents only 2% of total body weight, it uses ∼20% of the body’s total glucose to sustain neuronal survival and neurotransmission (12). Periods of heightened neuronal activity are closely tied to increases in regional glucose consumption and aerobic glycolysis (13). According to the astrocyte-neuron lactate shuttle (ANLS) hypothesis, lactate rather than glucose is preferentially used as a fuel source to support glutamatergic activity in neurons (14, 15). Excitatory neurotransmission stimulates glucose uptake across the blood brain barrier and astrocytic glycolysis. Astrocytes produce lactate as a metabolic end product of glycolysis and shuttle it to neurons as fuel for mitochondrial respiration and ATP generation (16). Therefore, lactate levels within the interstitial fluid (ISF) of the brain often serve as a proxy for excitatory neuronal activity (15-17). ISF lactate also has a diurnal rhythm (18), which follows neuronal firing patterns across sleep-wake periods. Neuronal firing rates are increased during wake as compared to NREM (19, 20). Therefore, lactate rises acutely upon a transition to wake and falls following transitions to NREM (21-24). Because of this, lactate can also be considered a biomarker for sleep-wake transitions (25).

While the relationships between cerebral metabolism and neuronal activity are well described during euglycemia, it is unclear how simple changes in energy availability, such as bouts of hyperglycemia or hypoglycemia, can alter the relationship between glucose, lactate, and sleep. Peripheral hyperglycemia is sufficient to increase ISF glucose and ISF lactate in mice (26). Additionally, recurrent hypoglycemia increases lactate transport into the type-1-diabetes brain. Together, this indicates that modulating systemic glucose can alter lactate dynamics in the brain which could then, in turn, could regulate sleep (27, 28). As increased lactate is associated with increased neuronal activity and waking behaviors, we hypothesized that acute bouts of hyper- and hypoglycemia are sufficient to alter sleep-wake states due to changes in ISF lactate. To evaluate this hypothesis, we paired amperometric biosensors detecting ISF glucose and lactate with EEG/EMG recordings in unanesthetized, unrestrained wildtype mice. First, we established a metabolic signature of sleep/wake cycles using acute fluctuations in ISF glucose and lactate, diurnal oscillations, and power spectral analyses during euglycemia. Next, we performed acute glycemic challenges, representing bouts of hyper- and hypoglycemia, to test how fluctuations in blood glucose levels relate to changes in ISF glucose, ISF lactate, and sleep/wake patterns on a sub-second timescale. During euglycemia, our results show a consistent anti-correlated relationship between ISF glucose and lactate across sleep states, but coherence and phase shift depending on the frequency of their oscillation. Further, we demonstrate specific changes in ISF glucose and lactate exist within different frequency bands during sleep-wake states. Most notably, ISF lactate, not ISF glucose, is closely associated with changes in the EEG power spectrum during wake. These relationships are absent during NREM periods. During hyper- and hypoglycemia, we defined how changes in blood glucose levels uncouple the typical relationship between glucose and lactate. Both hyper- and hypoglycemia increased ISF lactate levels and decreased NREM sleep, decreased delta band activity, increased theta band activity, and increased EEG coherence within delta, theta, sigma, and high gamma bands. Moreover, systemic injections of lactate are sufficient to drive increased ISF lactate and disrupt sleep. Hyper- and hypoglycemia also disrupt the typical relationships between ISF glucose, lactate, and specific EEG frequency bands. Relationships between ISF lactate and different EEG spectral bands typically found during wake emerge during NREM sleep in response to hyper- and/or hypoglycemia. More specifically, alterations in peripheral glucose metabolism stimulates aberrant, high gamma activity typically absent during NREM, leading to disrupted sleep. Together, our data suggests that changes in blood glucose levels are sufficient to uncouple the metabolic signature of sleep-wake states by altering the dynamics of ISF lactate, which appears to subsequently disrupt sleep.

## Results

### The metabolic signature of sleep-wake behaviors during euglycemia

First, we characterized the metabolism of sleep-wake and diurnal cycles during euglycemia. Cosinor analysis of the diurnal rhythm of ISF glucose and lactate showed both metabolites are increased during the dark period and decreased during light period (Fig 2A-B, p<0.0001). However, only ISF lactate shows a difference across sleep-wake states independent of light/dark periods, with ISF lactate elevated during wake (Fig 2B, p<0.05). During a NREM-to-wake transition, ISF lactate rises 1.5 minutes post-transition (Fig 2C, p<0.001) while ISF glucose rises above baseline levels after 6.5 minutes (p<0.01). During a wake-to-NREM transition, ISF lactate decreases below baseline after 40 seconds (p<0.0001) while glucose increases above baseline 50 seconds post transition (p<0.0001). Cross-correlation analyses revealed that ISF glucose and lactate are largely anti-correlated during both wake and NREM (Fig 2D). We performed multiple spectral analyses to examine the frequency specificity of this relationship. We found that both glucose and lactate follow a roughly 1/f pattern, although lactate has slightly greater power at higher frequencies relative to glucose (p<0.0001). Glucose and lactate are generally coherent across the range of frequencies examined; however, this coherence is reduced specifically around 0.01 Hz (Fig 2F). Notably, we found that this local drop in coherence corresponds with a shift from glucose and lactate being out of phase at frequencies <0.01 Hz to in phase at frequencies >0.01 Hz (Fig 2G). Thus, the anticorrelated nature of glucose and lactate is restricted to frequencies <0.01 Hz. The physiological significance of this observation is unclear at present; nonetheless, lower frequencies contribute substantially more variance to the observed signals, thus consistent with the predominantly anticorrelated appearance of the time series (Fig. 2D). Taken together, these data show that while both glucose and lactate fluctuate across the diurnal cycle, ISF lactate is more strongly associated with activity during wake as well as the transitions in the sleep-wake state. ISF glucose and lactate also have defined patterns across sleep-wake transition periods. Further, the temporal relationship between glucose and lactate is largely invariant to sleep state, although it is frequency-dependent. Together, these results demonstrate a clear metabolic signature of the sleep-wake system that exists under euglycemic conditions and confirm the importance of ISF lactate for wake-related transitions and behaviors.

**Figure 1.**
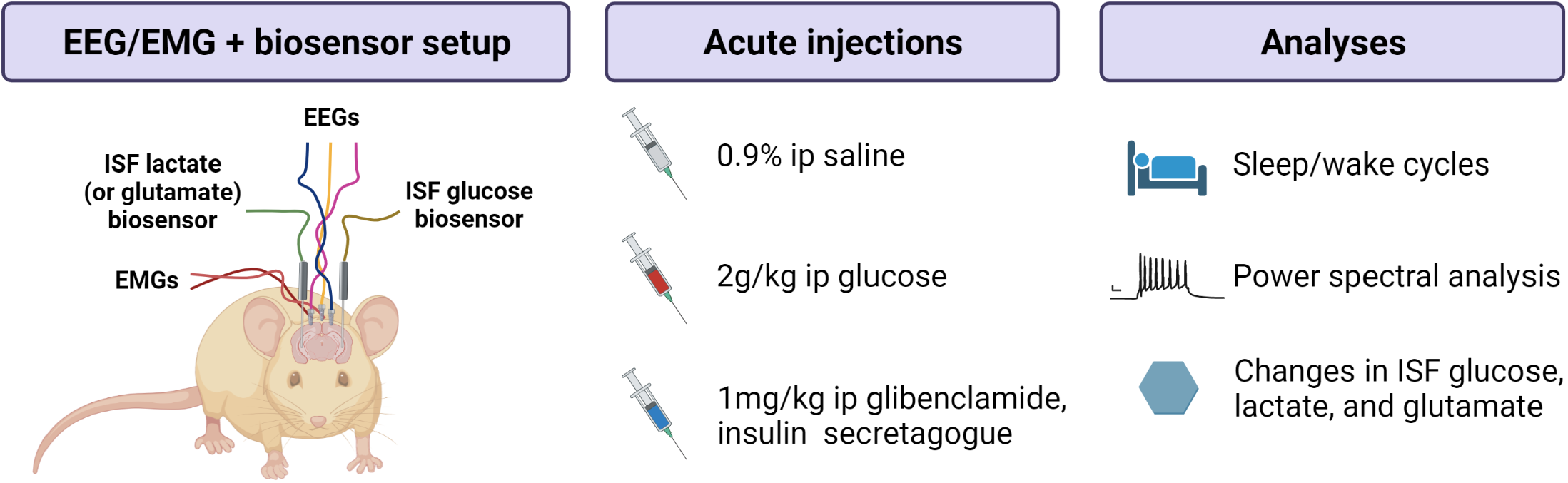
Experimental design and effect of acute hyper- and hypo-glycemia on peripheral metabolism. Representation of biosensor and EEG/EMG set-up and acute injection analysis workflow.

### Acute hyper- and hypoglycemia disrupt the typical relationship between ISF glucose and lactate

Next, we explored whether changes in blood glucose were sufficient to alter glucose and lactate levels in both the periphery and hippocampal ISF. IP glucose injections, mimicking acute hyperglycemia, raised blood glucose levels 167.7 ± 4.8%, with a peak at 15 minutes post injection. Similarly, IP glibenclamide injections, an insulin secretagogue that induces hypoglycemia, lowered blood glucose levels by 53.9 ± 2.5% (Fig 3A). This effect was observed at 30 minutes post-injection and persisted for the hour sampling period. Interesting, neither acute hyper-or hypoglycemia altered blood lactate levels (Fig 3A). Acute episodes of hyper- and hypoglycemia rapidly changed glucose and lactate levels in the brain’s ISF by 10-15 minutes post-injection. Glucose injections caused an 83.7± 7.7% increase in ISF glucose over a baseline saline response, with a peak ∼25 minutes post injection (Fig 3B-C, p<0.001). There was also a 19.2 ± 6.3% increase in ISF lactate, peaking ∼25 minutes post injection (Fig 3B-C; p<0.05). Glibenclamide injections caused a 47.7 ± 3.1% decrease in ISF glucose with a max decrease ∼40 minutes post injection (Fig 3B-C, p<0.001). ISF lactate increased 17.3 ± 5.4% over baseline, with peak occurring ∼20 minutes post injection (Fig 3B-C; p<0.05), representing an uncoupling of the typical relationship between glucose and lactate. There were no changes in ISF glutamate levels following either injection compared to saline response (Supp. Fig 1A). The increases in ISF glucose and lactate during hyperglycemia were strongly correlated (Fig 3D; p<0.01), while the increase in ISF glutamate and glucose trended towards significance. Increases in ISF glutamate and lactate were also correlated, illustrating the relationship between ISF lactate and excitatory neurotransmission (Supp. Fig 1B, p<0.001). Acute hypoglycemia caused a negative correlation between ISF lactate and glucose (Fig 3D; p<0.01) and ISF glutamate and glucose (Supp. Fig 1B, p<0.001). These data demonstrate peripheral hyper- and hypoglycemia can modulate ISF glucose levels in a similar manner to changes observed in the peripheral blood glucose. However, ISF lactate is modulated independent of any changes in blood lactate levels, indicating changes in ISF lactate are in response to the cerebral glucose uptake and metabolism.

**Figure 2.**
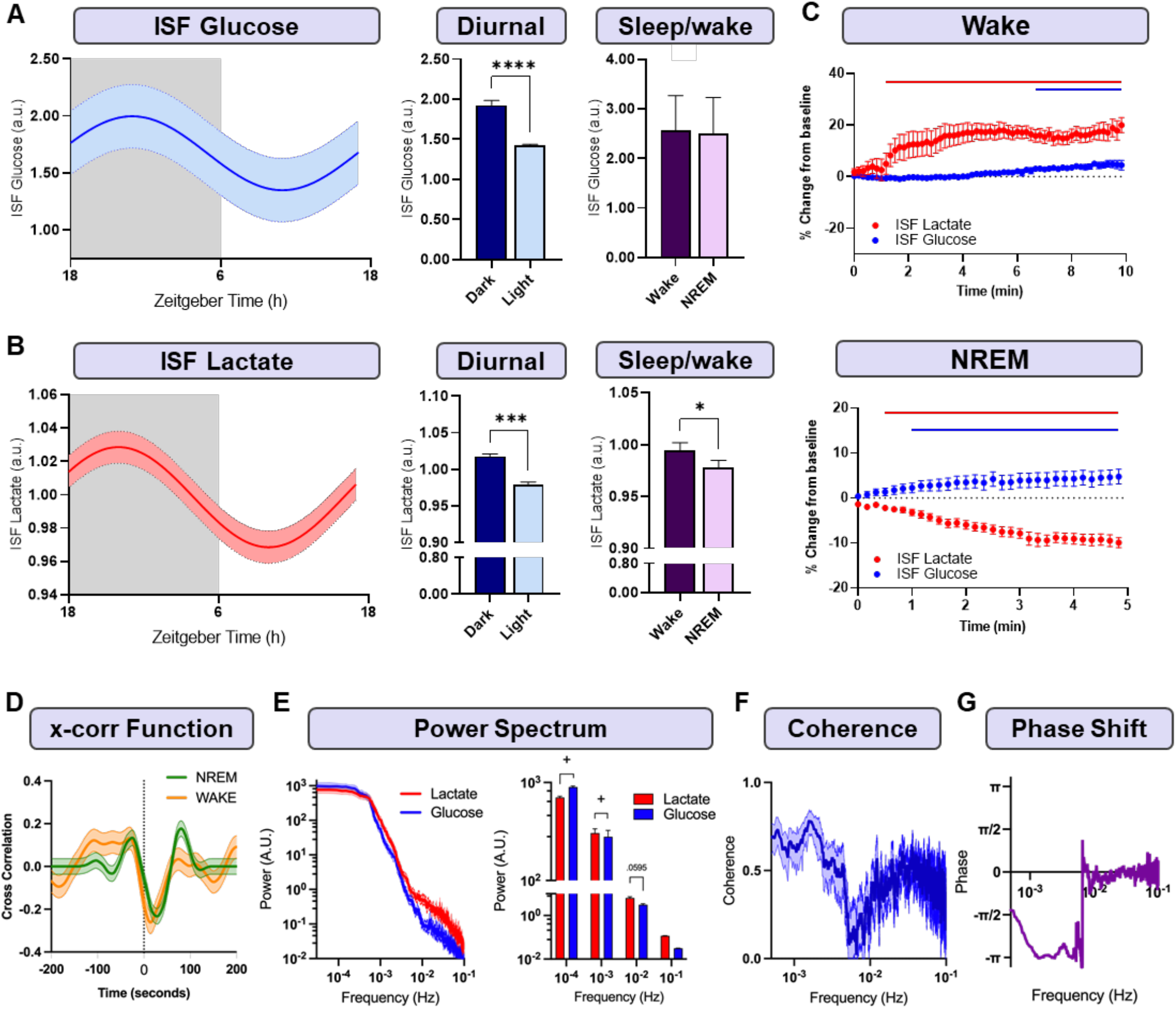
Relationship between glucose and lactate oscillations across diurnal and sleep/wake cycles. (**A-B**) Diurnal rhythm of ISF glucose and lactate as shown by cosinor curves. Both ISF metabolites increased during the dark period (p<0.001). ISF lactate, but not ISF glucose, is elevated during wake compared to NREM (p<0.05). (**C**) Sleep state also influences ISF glucose and lactate across sleep-wake transitions. In a NREM-to-wake transition, ISF lactate increases immediately above baseline, followed by an increase in glucose. During a wake-to-NREM transition, ISF lactate decreases immediately while ISF glucose increases. Red line indicates significant change from baseline in ISF lactate. Blue line indicates significant change in ISF glucose. (**D**) Cross correlation function demonstrates anti-correlated relationship between acute fluctuations in ISF glucose and lactate that is independent of sleep state. Glucose decreases precede lactate increases by ∼10 seconds. (**E**) Power spectral analysis of glucose and lactate reveal a ∼1/f pattern for both metabolites, but with glucose having relatively greater power at lower frequencies in comparison to lactate (p<0.0001, p<0.05). (**F-G**) Coherence analysis shows a localized drop in coherence between glucose and lactate around 0.01 Hz that corresponds with a shift from out of phase (<0.01 Hz) to in phase (>0.01 Hz). Data reported as means ± SEM. n = 5-8 mice/condition ^*^p<0.05, ^#^p<0.01, ^^^p<0.001, ^+^p<0.0001 using 1-way or 2-way ANOVAs and Dunnett’s post-hoc tests.

**Figure 3.**
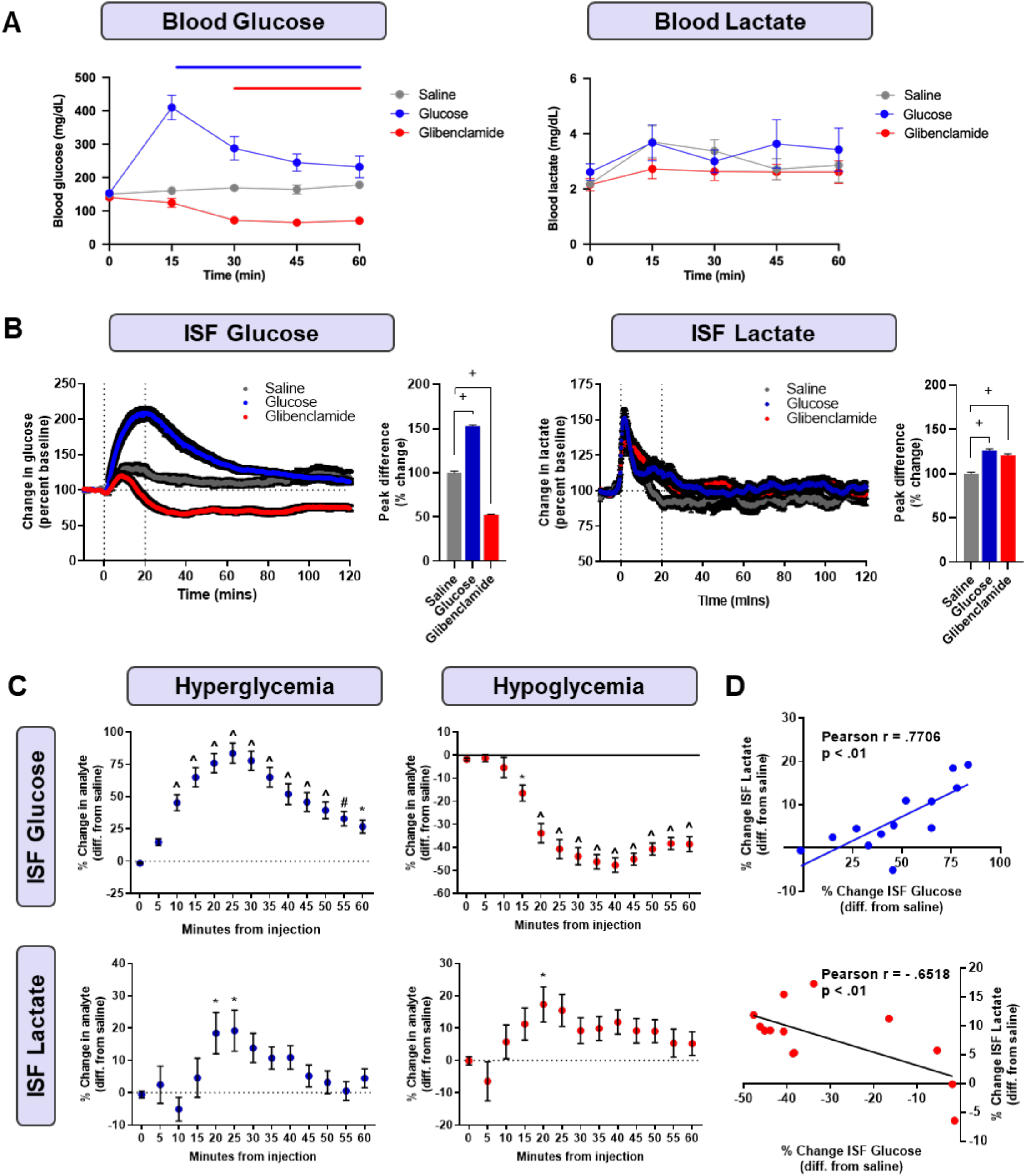
Acute hyper- and hypo-glycemia alter ISF glucose and lactate levels in the ISF but not periphery. (**A**) Acute hyperglycemia via IP glucose injection (2g/kg) increased peripheral blood glucose levels, with the peak concentration 15 minutes post-injection (167.7 ± 4.8%). Acute hypoglycemia via IP. glibenclamide injection (1mg/kg), a sulfonylurea that stimulates insulin release, decreased blood glucose, with a trough 45 minutes post-injection (-53.9 ± 2.5%). Blood lactate levels were not changed from baseline saline response. *Blue line = p<0*.*05 for glucose injections compared to saline, red line = p<0*.*05 for glibenclamide injections compared to saline*. (**B-C**) Acute hyperglycemia increased ISF glucose. Peak ISF glucose occurred 25 minutes post-injection (83.7 ± 7.7% increase over baseline). Acute hypoglycemia decreased ISF glucose, with a trough 40 minutes post-injection (-47.7 ± 3.1%). Acute hyper- and hypo-glycemia increased ISF lactate, uncoupling of the glucose-lactate relationship observed during euglycemia. Peak ISF lactate occurred 25 minutes post-glucose injection (19.2 ± 6.3%) and 20 minutes post-glibenclamide injection (17.3 ± 5.4%). (**D**) The rises in ISF glucose and lactate during acute hyperglycemia were positively correlated (*r =* 0.7706. p = 0.0021). During acute hypoglycemia, ISF glucose was negatively correlated with ISF lactate (*r =* -0.6518. p = 0.0158). Data are reported as means ± SEM. *n* = 10/peripheral experiment, 20 per ISF glucose biosensor and 12 per ISF lactate biosensor.^*\**^*p<0*.*05*, ^*#*^*p<0*.*01, ^p<0*.*001*, ^+^p< 0.0001 using 1-way or 2-way ANOVAs and Dunnett’s post-hoc tests.

### Acute hyper- and hypoglycemia increase time spent awake by changes in ISF lactate

Next, we explored if uncoupling glucose and lactate disrupts sleep-wake patterns. Both hyper- and hypo-glycemic conditions increased time spent awake and decreased time spent in NREM two hours post-injection (Fig 4A-B; p<0.01). Time spent awake following glucose injection was correlated with increased ISF lactate (Fig 4D; p <0.01) and ISF glucose (Fig 4D; p<0.001), and trended towards increased ISF glutamate (Supp. Fig 1B). Time spent awake post glibenclamide injection was positively correlated with ISF lactate (Fig 4E; p < 0.001) and ISF glutamate (Supp. Fig 1C; p< 0.01), but negatively correlated with ISF glucose (Fig 4E; p<0.0001). These studies demonstrate that acute alterations in blood glucose levels perturb sleep by increasing ISF lactate levels.

**Figure 4.**
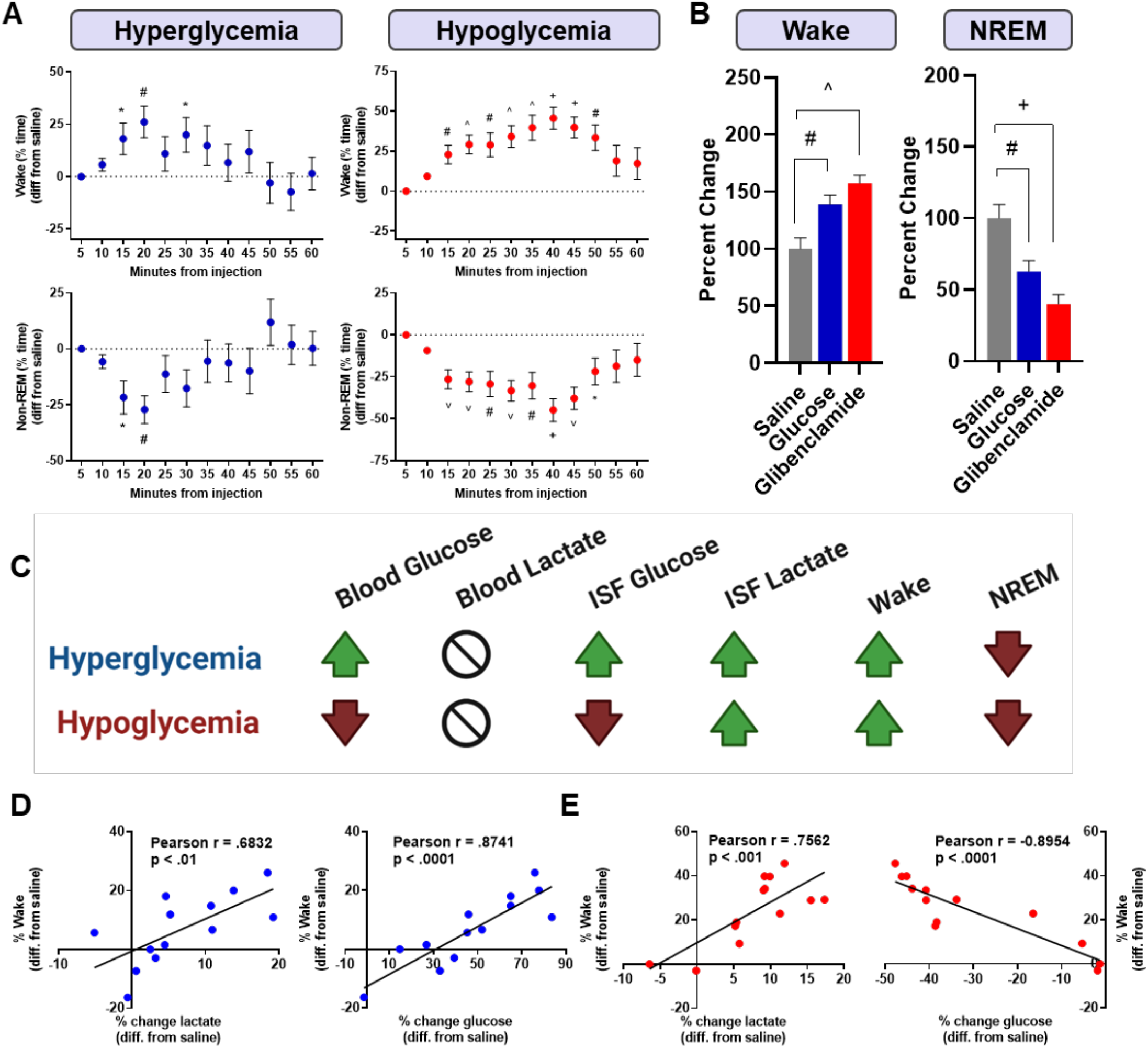
Acute hyper- and hypoglycemia is sufficient to increase wake and decrease NREM. **(A-B)** Acute hyper- and hypoglycemia increased time awake (hyperglycemia: 39.1 ± 8.0% increase over saline, hypoglycemia: 57.3 ± 7.2%) and decreased NREM (hyperglycemia: -37.3 ± 9.7%, hypoglycemia: -60 ± 6.7%) (C) Summary of changes associated with glycemic challenges (D) Increased wake during hyperglycemia was positively correlated with increased ISF glucose *(r* = 0.8741, p < 0.0001) and ISF lactate *(r* = 0.6832 p < 0.01). (E) Time awake during hypoglycemia was negatively correlated with ISF glucose (*r =* -0.8954, p <0.0001), but positively correlated with ISF lactate (*r =* 0.7562. p < 0.001). Data are reported as mean ± SEM. *n* = 11-12 mice/group. ^*\**^ *p<0*.*05*, ^*#*^*p<0*.*01*, ^*^*^*p<0*.*001*, ^*+*^*p<0*.*0001* using 1-way ANOVAs and Dunnett’s post-hoc tests.

To demonstrate that changes in ISF lactate are sufficient to disrupt sleep, we performed systemic IP injections of lactate and investigated the effects on ISF lactate, time spent in NREM, and qEEG (Supp. Fig 2). Systemic lactate injections increase ISF lactate, stimulate wake, and decrease NREM. There was also increased delta activity suggesting an increased propensity for sleep (Supp. Fig 2).

### Peripheral glycemic state impacts temporal relationship between ISF lactate and glucose

Next, we investigated how fluctuations in ISF glucose and lactate are temporally related as a function of both sleep and glycemia using a cross correlation analysis. During euglycemia, ISF lactate and glucose are anti-correlated. During hyperglycemia and hypoglycemia across both wake and NREM, this relationship is altered as indicated by the shifts in cross correlation traces (Fig 5A). During hyperglycemia in wake, ISF glucose and lactate display both a strong anti-correlated and correlated relationship which is a deviation from typical wake-euglycemia. During hypoglycemia in wake, the anti-correlated relationship between ISF glucose and lactate is lost and replace by a correlated relationship of decreased magnitude. During NREM, the strong anti-correlated relationship between glucose and lactate is lost both hyper- and hypo-glycemia and replaced by correlated relationship between ISF glucose and lactate. This relationship is particularly strong during hypoglycemia. A power spectral analysis of ISF glucose and lactate oscillations shows both hyper- and hypoglycemia increase power of ISF lactate (p<0.0001, p<0.01; Fig 5B) and glucose (p<0.05, p<0.01; Fig 5B) at low frequencies. Conversely, glycemic challenges cause decreased power at higher oscillation frequencies, again in both ISF lactate (p<0.05, 0<0.0001, Fig 5B) and ISF glucose (p<0.001, p<0.01, Fig 5B). Therefore, hyper- and hypoglycemia are sufficient to disrupt the temporal relationship between ISF glucose and lactate.

**Figure 5.**
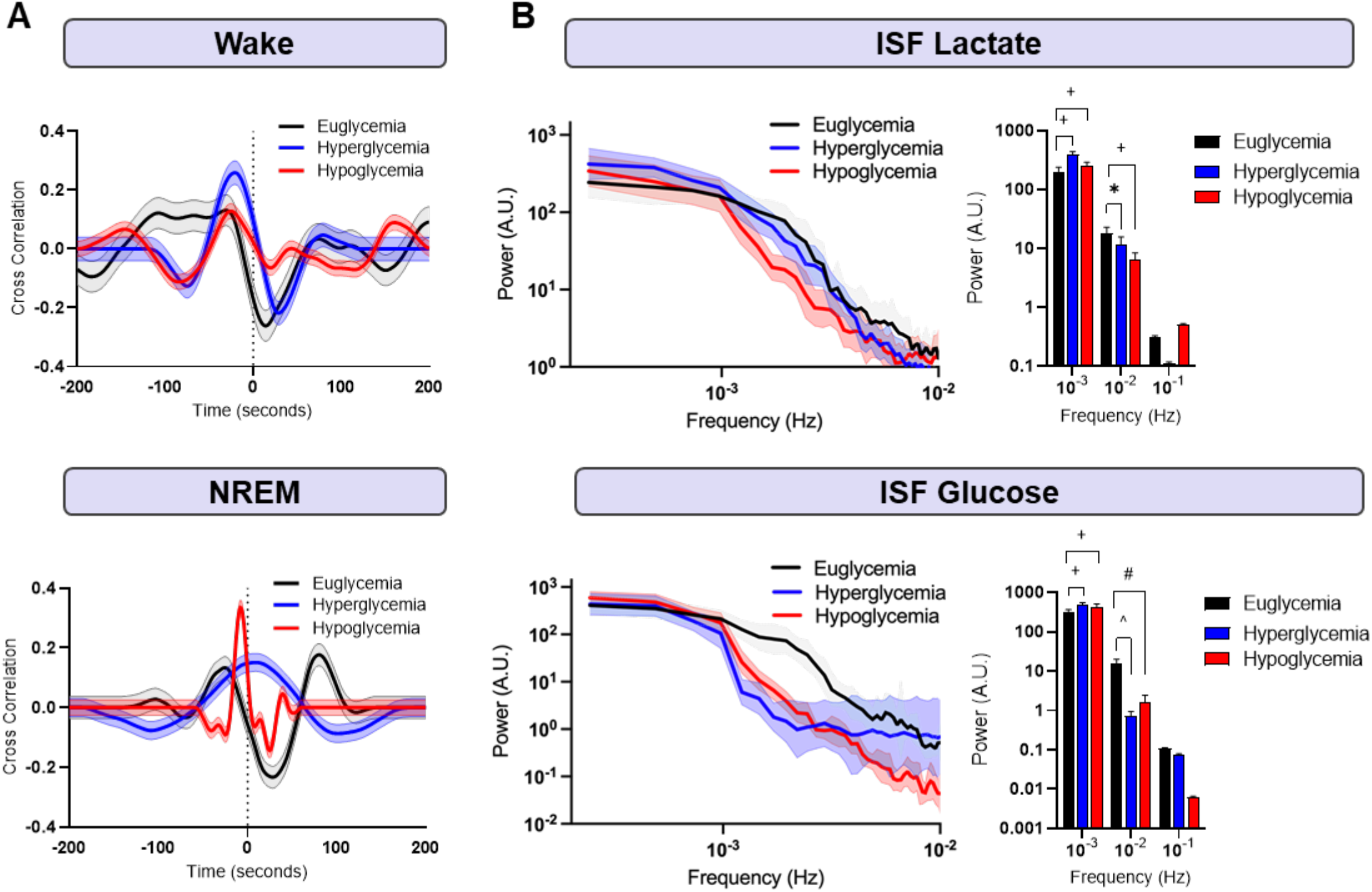
Hyper- and hypoglycemia disrupt relationship between glucose and lactate across sleep states. **(A)** Across sleep states, ISF glucose and lactate are anti-correlated during euglycemia. This relationship is altered by hyper- and hypo-glycemia. (**B**) Power spectral analysis of glucose and lactate shows glycemic challenges disrupt the power distribution of analytes independent of sleep state. Both hyper- and hypoglycemia increased power of ISF lactate at low frequencies (p<0.0001, p<0.01, respectively). For ISF glucose, a dramatic shift in the power spectrum is clear, with hyper- and hypoglycemia increasing power of ISF glucose in low frequencies compared to euglycemic conditions (p<0.05, p<0.01, respectively). ^*^*p<0*.*05*, ^#^*p<0*.*01*, ^^^*p<0*.*001*, using two-way ANOVAs with multiple comparisons corrections. *n* = 5-6 mice/group.

### Hyper- and hypoglycemia eliminate the metabolic signature of sleep to wake transitions

Both glucose and lactate have well-characterized patterns across sleep-wake transitions, where lactate rises at the onset of wake and drops with NREM (24, 25). These relationships, however, are only understood during euglycemic conditions. Therefore, we investigated how hyper-or hypoglycemia alters the metabolic phenotype of a transition period between sleep and wake. Acute hyper- and hypoglycemia blunt the increase in lactate during a NREM-to-wake transition, with no increase above baseline (Fig 6D, p<0.001). During hyperglycemia, ISF glucose does not change from baseline, in contrast to the increase seen during euglycemia (Fig 6D, p<0.001). In hypoglycemia, ISF glucose decreases from baseline from 1-5 minutes following a transition to wake (Fig 6B; p<0.05) and never rises above baseline, again in contrast to euglycemia (Fig 6D, p<0.0001). Conversely, in a wake-to-NREM transition, the decrease in ISF lactate is unaffected by hyper- and hypoglycemia (Fig 6C). ISF glucose does not change from baseline in hyperglycemia, again resulting in a decreased max change from baseline compared to euglycemia (Fig 6D, p<0.05). During hypoglycemia, ISF glucose increases above baseline (Fig 6D, p<0.001) in a similar manner to euglycemia. These analyses show peripheral changes in blood glucose levels alter the metabolic patterns of sleep-wake transitions, with the largest impact occurring during NREM-to-wake transitions where the increase in ISF lactate is blunted which could have implications for wake-associated behaviors.

**Figure 6.**
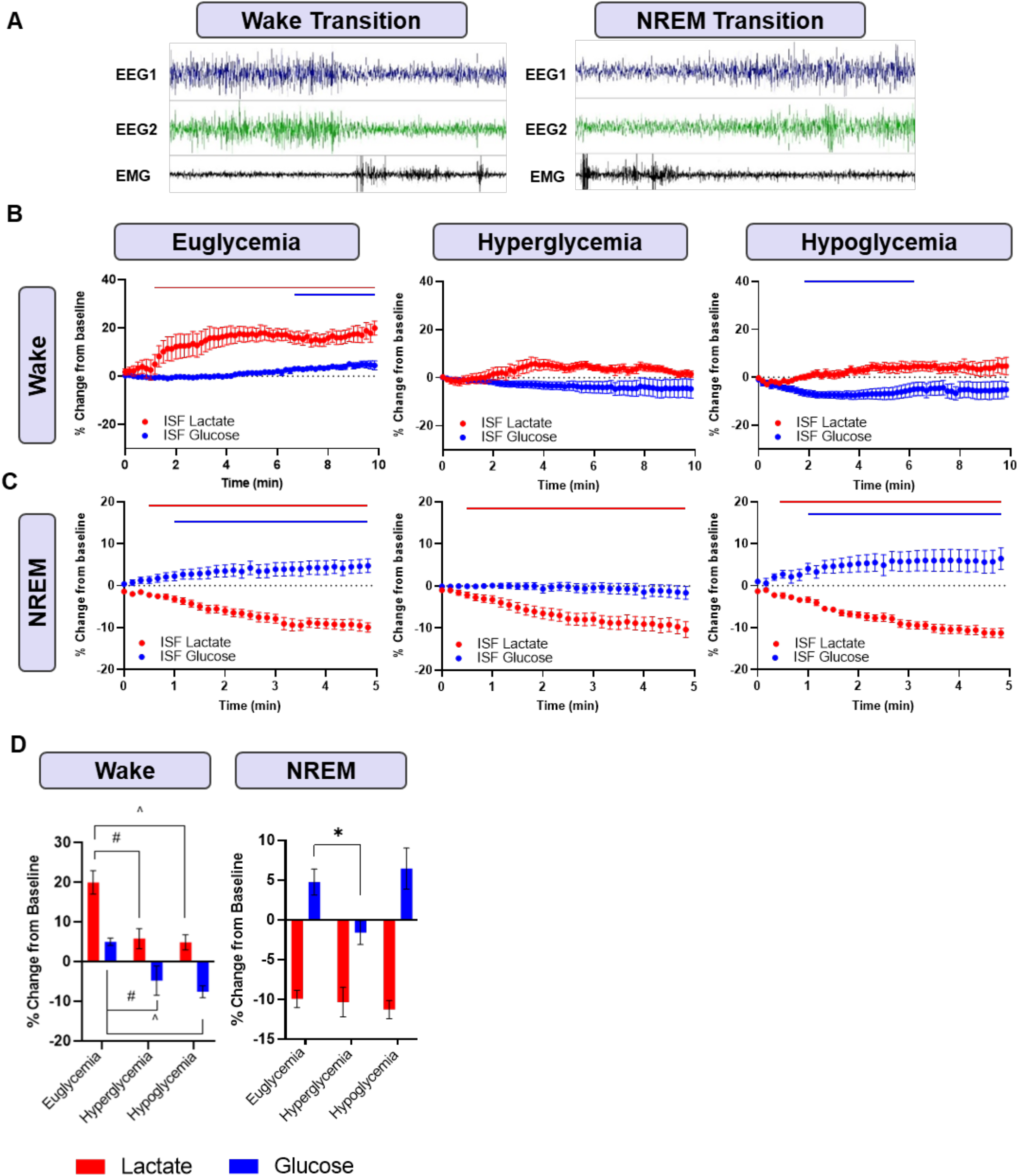
ISF lactate increase during NREM-to-wake transition is abolished during hyper- and hypo-glycemia. **(A)** Representative EEGs and EMGs during transitions. **(B)** In euglycemia, ISF lactate immediately increases following a NREM-to-wake transition (20 ± 3.0 %). ISF glucose shows an increase above baseline ∼6 minutes post transition (5 ± 0.9%). Hyperglycemia blunts ISF lactate and glucose increases. In hypoglycemia, ISF glucose decreases initially following a transition to wake (-7.6± 1.6%), while ISF lactate does not change. **(C)** During a wake-to-NREM transition in euglycemia, ISF lactate decreases immediately (-10 ± 1.1%), while glucose increases (4.7± 1.6%). During hyperglycemia, ISF lactate decreases (-10.4± 1.9%), but ISF glucose remains unchanged. In hypoglycemia, ISF lactate decreases (-11.3 ± 1.1%). (D) Across NREM-to wake transitions, the maximal change in lactate is decreased with glycemic challenges. ISF glucose has a positive max change during euglycemia, but negative during both glycemic challenges. During wake-to-NREM transitions, ISF glucose increases during euglycemia and decreases during hyperglycemia. Data reported as means ± SEM. n = 6 mice/group. *Red line* = *p<0.05* for lactate, *blue line* = *p*<0.05 for glucose. \**p*<0.05. #p<0.01, “*p<0.001*, +*p<0.0001*, using a 1-way ANOVA and Dunnett’s post-hoc tests.

### Hyper- and hypoglycemia alter spectral power density and relative power characteristics

Given the changes in overall sleep time during glycemic challenges, we next investigated how hyper- and hypoglycemia impacts power spectral characteristics. Spectral power was binned based on defined frequency bands including delta (0.5-4 Hz), theta (4-8 Hz), alpha (8-13 Hz), sigma (12-15 Hz), beta (13-30 Hz), gamma (30-50 Hz), and high gamma (50-100 Hz). In saline conditions, there was a typical state-dependent distribution with peaks across delta and theta frequencies during wake and high delta power during NREM. Hyper- and hypo-glycemic challenges reduced power in low frequencies (1-5 Hz), during both wake and NREM (Fig 7A; p<0.05). Next, shifts in relative power bands were used to evaluate how changes in total power are distributed into pre-defined frequency bands. During wake, relative delta power, a marker of homeostatic sleep drive, decreased during hyper- and hypoglycemia (Fig 7B; p<0.01, p<0.001). Hypoglycemia increased relative theta power (Fig 7B; p<0.05), indicating increased sleep propensity (30). During NREM, hyperglycemia and hypoglycemia decreased relative delta power (Fig 7B; p<0.001, p<0.01). Both injections also increased theta power (Fig 7B; p<0.05). The ratio of relative theta to gamma power, which is associated with cognitive decline and memory impairment (31, 32), increased during hypoglycemia supporting existing literature that hypoglycemia is negatively associated with cognition (33-36) (Fig 7C; p<0.05).

**Figure 7.**
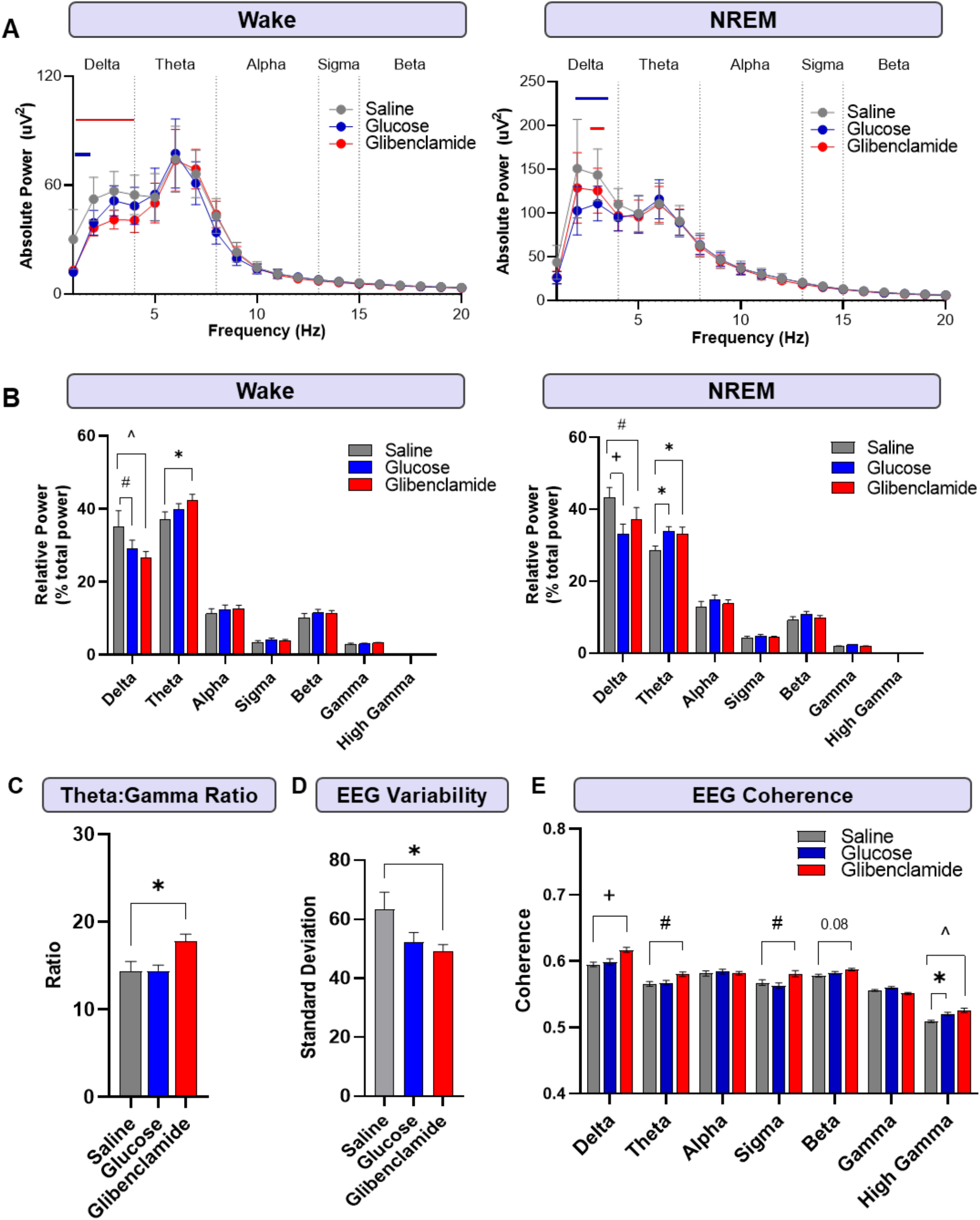
Hyper- and hypo-glycemia decrease sleep and disrupt EEG characteristics. **(A)** Absolute power was reduced in low frequencies (1-4 Hz) during hyper- and hypo-glycemia across both wake and NREM. Blue line = p<0.05 for glucose injection, red line = p<0.05 for glibenclamide injection. (**B**) Both injections decreased relative delta power during wake (-17.4± 1.8% and -24.5± 1.8%, respectively). Glibenclamide increased relative theta power (14.0± 1.4%). During NREM, both injections decreased relative delta power (-23.3 ± 1.7% and 14.2 ± 1.8%, respectively). Both injections also increased relative theta power (18.6 ± 1.1% and 15.6 ± 1.2%). (**C**) The ratio of relative theta: gamma power increased during hypoglycemia during wake (23.4± 1.0% compared to saline; 23.1± 0.9% compared to glucose). (**D**) EEG variability, as calculated by the standard deviation of total EEG signal fluctuations, decreased during hypoglycemia (-24.7± 1.9) (E) Coherence between two EEG leads increased during hypoglycemia within delta (3.7± 0.03% compared to saline injection), theta (1.5± 0.03%), sigma (2.6± 0.06%), and high gamma bands (3.3± 0.02%). During hyperglycemia, high gamma coherence increased compared to saline (2.2 ± 0.05%). Data are reported as means ± SEM. n = 10-11 mice/group. ^*\**^ *p<0*.*05*, ^*#*^*p<0*.*01*, ^*^*^*p<0*.*001*, ^*+*^*p<0*.*0001* using 2-way ANOVA with Tukey’s multiple comparisons correction for post hoc analysis and one-way ANOVA with Dunnett’s post-hoc test.

### Hyper- and hypoglycemia decreases EEG power while increasing EEG coherence

Given changes observed in spectral power, we asked whether overall EEG amplitude or coherence between EEG leads was affected by hyper- and hypoglycemia. EEG amplitude, a marker of EEG power, decreased during hypoglycemia compared to the saline response (Fig 7D; p<0.05). EEG coherence measures the level of synchrony between two distinct brain regions and is often used to show functional connectivity of two distinct regions; however, too much coherence is associated with increased seizure activity especially in high frequency bands. Coherence between parietal and frontal EEG leads was increased during hypoglycemia within the delta (p<0.0001), theta (p<0.01), sigma (p<0.05), and high gamma bands (p<0.01) compared the saline response. In hyperglycemia, only high gamma coherence increased compared to the saline condition, which is typically associated with seizure activity (Fig 7E; p<0.05). Overall, these changes demonstrate EEG activity is altered by hyper- and hypoglycemia, although hypoglycemia seems to have a more pronounced effect on EEG amplitude and coherence.

### Hyper- and hypoglycemia alter associations between spectral power bands on ISF metabolites

Next, we explored whether activity within certain spectral power bands is related to fluctuations in ISF glucose and lactate, supporting the idea that glucose is recruited to specific brain regions in an activity-dependent manner. To do this, we used a general linear model across both wake and NREM bouts in each glycemic condition. In general, ISF lactate was associated with changes in activity across multiple frequency bands during wake. ISF lactate was positively associated with sigma (Fig 8A; p<0.05) and high gamma (p<0.05) band activity and negatively associated with delta (p<0.05), alpha (p<0.001), and gamma (p<0.01) band activity. Across wake in euglycemia, changes in alpha power are positively correlated with changes in ISF glucose (Fig 8A; p<0.05). Hyper- and hypoglycemia ablated these relationships, demonstrating the metabolic signature of the power spectrum is lost with metabolic dysregulation. In particular, all associations were lost during hyperglycemia, with the exception of ISF lactate which showed a positive association with alpha (p<0.05). During hypoglycemia, no correlations exist. A different set of relationships emerged during NREM, where few relationships existed between NREM and ISF lactate or glucose during euglycemia. Theta band activity was positively associated with both ISF glucose (Fig 8B; p<0.05) and ISF lactate (p<0.05), while high gamma activity was positively associated with ISF lactate (p<0.05). Hyper- and hypoglycemia increased the number of relationships between spectral power bands and ISF lactate or glucose. During hyperglycemia, ISF glucose was positively associated with theta (p<0.05) and high gamma (p<0.01) activity and negatively associated with changes in delta (p<0.05) and sigma (p<0.05) activity. ISF lactate showed changes within the same bands, but the relationships were opposite; delta (p<0.01) and sigma (p<0.01) band were positively associated, while theta (p<0.05) and high gamma (p<0.01) were negatively associated. During hypoglycemia, ISF glucose had no associations with different power spectral bands, while several new relationships emerged for ISF lactate that were absent during euglycemia. Theta (p<0.01), alpha (p<0.05), sigma (p<0.01), and high gamma (p<0.05) activity were all positively associated with changes in ISF lactate while beta (p<0.001) activity was negatively associated. The increased association with ISF lactate and NREM during hypoglycemia resembled patterns seen during wake, suggesting hypoglycemia triggers changes in ISF lactate that disrupts sleep. Together, this demonstrates that wake possesses a unique set of relationships with spectral power bands and ISF lactate. These associations are ablated during hyper- and hypoglycemia. Conversely, in NREM, hyper- and hypoglycemia produce new relationships between frequency bands and ISF lactate or glucose that are not traditionally present during euglycemia. In several cases, the new relationships between power spectral bands, ISF lactate, and ISF glucose resembled wake rather than sleep.

**Figure 8.**
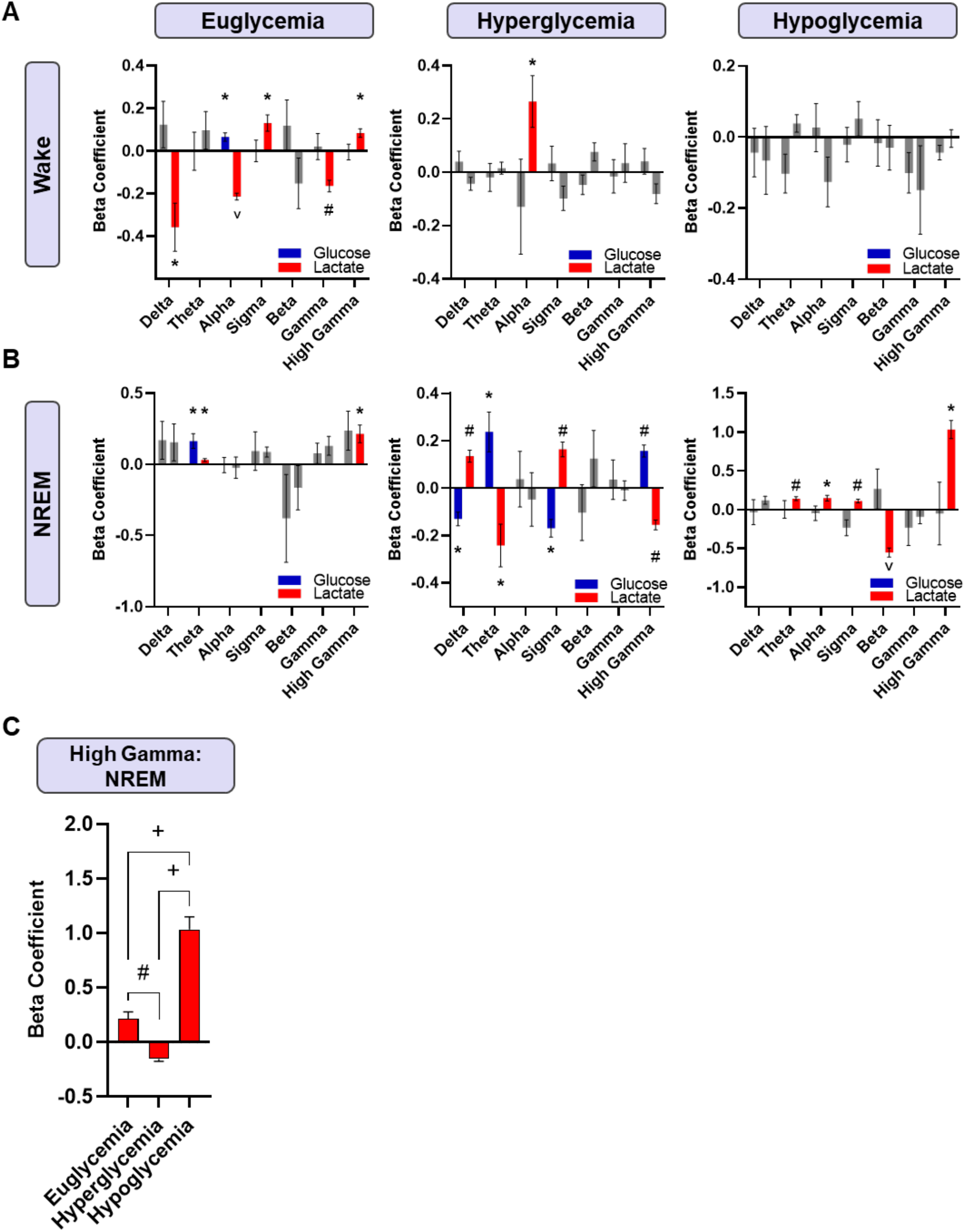
The metabolic signature of spectral band activity is lost with hyper- and hypoglycemia according to general linear model. **(A)** Changes in ISF glucose during euglycemic wake are positively associated with alpha band power. Changes in ISF lactate are positively associated with theta and high gamma and negatively associated with delta, alpha, and gamma activity. During a hyperglycemic challenge, the relationship with ISF glucose is lost, and ISF lactate is only positively associated with alpha activity. During hypoglycemia, no associations are significant. **(B)** During euglycemic NREM, changes in ISF glucose are positively associated with theta activity. ISF lactate is positively associated with theta and high gamma activity. In hyperglycemia, ISF glucose is negatively associated with delta and sigma activity and positively associated with theta and high gamma activity. ISF lactate is positively associated with delta and sigma activity and negatively associated with theta and high gamma activity, opposite associations as those seen with ISF glucose. No significant associations exist with ISF glucose during NREM in hypoglycemia. However, ISF lactate is positively associated with theta, sigma, alpha, and high gamma activity and negatively associated with beta activity. **(C)** During NREM, ISF lactate shows changes within high gamma band activity. In euglycemia a positive relationship exists; this relationship switches to negative during hyperglycemia and is exacerbated in hypoglycemia. Data are reported as means ± SEM. *n* = 10 mice/condition. \**p*<0.05, #*p<0.01*, \**p<0.001*, +*p<0.0001*. One sample t tests were used to determine significance of each beta coefficient from zero. One-way ANOVA was used to evaluate significance between glycemic states.

Interrogating high gamma power more directly, ISF lactate showed a positive relationship with high gamma in NREM during euglycemia. This was reversed to a negative association during hyperglycemia (Fig 8C, p<0.01). Interestingly, hypoglycemia strengthened the positive relationship between ISF lactate and high gamma compared to what was observed during euglycemia (p<0.0001). This demonstrates the opposing effects glycemic extremes can have on the relationship between patterns of neuronal activity and brain metabolites. Taken together, these changes indicate (1) hyper- and hypoglycemia cause a loss in associations during wake, (2) hyper- and hypoglycemia increase associations during NREM, and (3) spectral band activity is more associated with changes in ISF lactate compared to glucose.

## Discussion

The coupling of metabolic and neuronal activity is necessary for state-dependent brain function. Changes in brain activity rely on the availability and delivery of nutrients (e.g. glucose) via local blood flow changes in a process termed functional hyperemia (37). This process is necessary to accomplish a variety of tasks and behaviors during wake and these relationships fundamentally change to support sleep. When this process is perturbed, healthy brain function is impaired. The sleep-wake cycle is a master regulator of behaviors and couples oscillations in metabolic and neuronal activity over the diurnal cycle. Sleep is one of the most evolutionary conserved processes, highlighting the importance of coordinated fluctuations in metabolic and neuronal activity. While it is well established that perturbations in the relationship between metabolic and neuronal activity underlie many pathological conditions, there is little known about how intrinsic alterations in metabolism can feed forward to disrupt coordinated processes like sleep.

To date, the relationship between cerebral metabolism and neuronal activity is only described in euglycemic conditions. Fundamental principles, like the ANLS, describe how metabolism changes in response to increased glutamatergic activity. Like many other processes, it fails to consider how metabolic disruptions, like hyperglycemia or hypoglycemia, impact metabolic- and activity-dependent cooperation in the brain despite the fact that metabolic impairment is associated with a variety of adverse health outcomes, most notably in individuals with diabetes. Uncontrolled blood glucose levels are present in prediabetes and diabetes, while diabetes treatment with insulin or sulfonylureas often leads to episodes of hypoglycemia. Independent of other complications from T2D, acute fluctuations in blood glucose levels are considered a potential predictor of poor health (38-41) including increased rates of white matter hyperintensities (42), gray matter atrophy, cognitive decline (43), cardiovascular disease, and all-cause mortality (40). While increased metabolism and neuronal activity are associated with waking behaviors, few studies to date explored how modulating ISF glucose and lactate levels directly impact sleep. Therefore, the goals of these experiments were to: (1) define the metabolic signature of sleep-wake states relative to the relationships between ISF glucose and lactate and EEG activity, (2) identify how changes in peripheral blood glucose levels influences those relationships, and (3) establish peripheral hyper- and hypoglycemia as direct modifiers of sleep independent of other comorbidities like diabetes, sleep apnea, cardiovascular disease, or neurodegenerative diseases, like Alzheimer’s disease.

### The metabolic signature of sleep-wake

Since our unique experimental setup allows us to pair biosensors concurrently measuring hippocampal ISF levels of glucose and lactate with cortical EEG/EMG recordings, we are able to explore how changes in glucose and lactate relate to metabolic and neuronal activity patterns across sleep-wake periods. While we first established that a diurnal rhythm exists for both ISF glucose and lactate, only ISF lactate is elevated across bouts of wake as compared to NREM sleep independent of the light/dark period. This indicates that changes in ISF lactate, rather than glucose, are connected to waking behaviors. This notion is further supported by sleep-wake transitions, where ISF lactate levels increase immediately following a transition to wake and decrease directly following a transition to NREM. We also found oscillations in ISF lactate are coupled with ISF glutamate, suggesting lactate is related to excitatory neurotransmission in the brain during wake. According to the cross-correlation analysis, the strongest relationship that exists between glucose and lactate across sleep states is anti-correlated, suggesting that rises in glucose are temporally linked with drops with lactate. However, using power spectrum, coherence, and phase analyses, we found two more sophisticated patterns emerge. First, we observed that lactate exhibits relatively increased power at higher frequencies in comparison to glucose. Moreover, at lower frequencies (i.e., <0.01 Hz), glucose and lactate display high coherence but are out of phase (anti-correlated). As frequencies increase towards 0.01 Hz, there is a local drop in coherence which corresponds with a shift from this anti-phase pattern to an in-phase relationship at frequencies >0.01 Hz. This frequency-dependent change potentially reflects glucose shifting to in-phase with increased lactate production and arousal fluctuations at faster timescales (44) and supports the idea that multiple, dynamic relationships exist between ISF glucose and lactate across arousal states. Our GLM results demonstrate that changes in ISF lactate are associated with changes in different power spectral bands only during wake, further supporting the idea that ISF lactate is necessary for waking behaviors. These relationships were not present for ISF glucose during wake, nor were the relationships with ISF lactate and power spectral bands present during NREM under euglycemic conditions, suggesting ISF glucose may be more important for basal, homeostatic activity. Together, these data establish a metabolic signature of sleep-wake in which increases in ISF lactate are connected with wake associated behaviors, likely due to increased neuronal firing rates. It further suggests that the concentrations of ISF lactate must drop and the associations between ISF lactate and different frequency bands must be lost in order to enter in and maintain NREM sleep. Not only does this support fundamental principles such as the ANLS and neuronal activity during wake, but it also supports established work in pathological conditions like Alzheimer’s disease where elevations in ISF lactate are associated with increased neuronal activity, sleep impairment, and amyloid and tau pathology (16, 18, 21, 26, 45-47). Our data also demonstrate that systemic administration of lactate is sufficient to raise ISF lactate, disrupt sleep, and increase time spent awake (Supp. Fig 2), illustrating ISF lactate is a direct modifier of sleep/wake states. Ultimately, these results also have implications for other diseases by suggesting that any deviations from peripheral glucose homeostasis can alter sleep-wake patterns, offering a potential mechanism for sleep disruptions commonly reported in T2D (11), independent of other factors like obstructive sleep apnea, restless leg syndrome, or nocturia.

### Acute hyper- and hypoglycemia disrupts the traditional relationships between glucose and lactate to impair sleep

After establishing a metabolic signature of sleep-wake rhythms during euglycemic conditions, we explored how modulating blood glucose would influence those relationships. First, we found hyper- and hypoglycemia bidirectionally modulated blood and ISF glucose levels, without affecting blood lactate levels. This suggests that alterations in ISF lactate are due to the CNS metabolism of glucose following peripheral metabolic challenges. Both hyper- and hypoglycemia increased ISF lactate, indicating both glucose rich and glucose poor environments drive increased neuronal activity. Together, these findings add further nuanced detail to the ANLS. Under euglycemic conditions, there should be a direct relationship between glucose and lactate in the brain in response to neuronal activity (15, 16). However, the ANLS assumes on-demand metabolic availability and fails to account for changes in glucose availability independent of task-based behaviors. Here, we show hypoglycemia is sufficient to disrupt the relationship between ISF glucose and lactate and uncouple the expected relationship between cerebral metabolism and neuronal activity (16). Both hyper- and hypo-glycemic conditions have implications for shifts in sleep-wake patterns since elevations in ISF lactate increase neuronal activity, increase time spent awake, and decrease NREM sleep.

While this work provides additional support for the idea that elevations in ISF lactate are sufficient to disrupt NREM sleep, we further explored why changes in glucose homeostasis alter sleep-wake patterns. First, we found that modulating peripheral glucose availability impacted the temporal relationship between glucose and lactate in the brain. Across wake and NREM, both hyper- and hypoglycemia altered the anti-correlated relationship found in euglycemia. This shift could be indicative of changes in lactate utilization due to heightened neuronal activity associated with increased wake (15, 17). While increased wake is generally associated with increased metabolic demand (48), hyper- and hypoglycemia increased power in low frequencies of ISF glucose and lactate, suggesting altered glucose availability can slow glucose and lactate oscillatory behavior despite increased demand. Second, we explored how hyper- and hypo-glycemia impact EEG power spectral characteristics. Overall, hyper- and hypoglycemia decreased delta power, or slow wave activity (SWA), and increased theta power. During wake, this likely reflects a decreased propensity for sleep (30, 49), while decreased SWA in NREM indicates decreased sleep quality and a loss of the restorative aspects of NREM. The increased theta power is associated with sleepiness and worse cognitive performance during wake (30, 50, 51), and sleep deprivation during NREM (52). Further, the theta to gamma ratio was increased, which is consistent with cognitive and memory impairments (32, 53). Taken together, these results show a general shift away from low frequency activity towards neuronal activity patterns more consistent with wake. This likely impairs neuronal recruitment into large, synchronous slow waves, as reflected in the loss of delta power. Using general linear modeling, nearly all spectral band activity was associated with changes in ISF lactate during euglycemic wake. Hyper- and hypoglycemia ablate most of these relationships, indicating that glucose abundance or deprivation is sufficient to disrupt those connections. In some cases, new relationships arise. During hyperglycemia, a positive association between alpha power and ISF lactate emerges. Alpha power is generally associated with quiet wakefulness and disappears during task-oriented behaviors, indicating a lack of higher order cognitive processing during hyperglycemic wake. During NREM, there are considerably fewer relationships during euglycemia as compared to wake. However, hyper- and hypoglycemia cause an increase in the number of associations between spectral activity and ISF glucose or ISF lactate. This shift explains the decrease in NREM seen during glycemic challenges. If atypical associations are made between metabolic and neuronal activity (e.g. ISF lactate and frequency bands), new patterns of activity more consistent with waking behavior could emerge, causing disruptions in generating and maintaining sleep. For example, high gamma activity is predominately present in euglycemic wake. However, ISF lactate and high gamma are associated in NREM during hyper- and hypo-glycemia suggesting the presence of spurious high frequency activity. Lastly, we found decreased EEG variability and increased EEG coherence which suggests a loss of specificity of brain networks and potential activation of inappropriate networks within the sleep wake circuits. Because the sleep-wake switch is maintained by the balance of excitatory and inhibitory pathways (54), any shift in activity could cause instability of the switch. In fact, hyper- and hypo-glycemia ablated the rise in ISF lactate across the NREM-to-wake transition. This is a departure from the expected immediate increase in ISF lactate and trailing rise in ISF glucose seen in euglycemic conditions. Again, this offers another explanation for why changes in peripheral glucose metabolism are sufficient to disrupt sleep. Together, these analyses show peripheral glycemic states can change how neuronal activity patterns are related to fluctuations in glucose and lactate in a state dependent manner and offer an explanation for the observed NREM disruptions.

Our studies established metabolic signatures of sleep-wake states and demonstrated how ISF glucose and lactate interact on multiple levels to maintain normal sleep-wake rhythms. Peripheral hyper- and hypoglycemia are sufficient to uncouple the typical relationship between cerebral metabolism, neuronal activity, and sleep. This leads to sleep disruptions, seen in decreased overall NREM, altered spectral characteristics, diminished metabolic signatures during transitions to wake, and deleterious associations between spectral band activity and ISF metabolites. Taken together, we propose peripheral glycemic state as a major contributor to sleep-wake disruptions, which has implications for exacerbating T2D, as well as countless other health conditions, such as Alzheimer’s disease.

## Materials and Methods

### Mice

Male and female young (3-5 month) wildtype mice on a B6C3 background were used in all experiments. Mice were given food and water *ad libitum* and maintained on a 12:12 light/dark cycle until surgery. All procedures were carried out in accordance with an approved IACUC protocol from either Washington University School of Medicine or Wake Forest School of Medicine.

### Peripheral glucose and lactate measurements

Peripheral changes in glucose or lactate were measured using a glucometer (Contour, Bayer) or lactate meter (Lactate Plus, Nova Biomedical) at 15, 30, 45, and 60 minutes post-injection.

### Surgery

Mice were anesthetized with isoflurane and guide cannulas (MD-2255, BASi Research Products) were stereotaxically implanted bilaterally into the hippocampus (from bregma, A/P -3mm, M/L +/- 3mm, D/V -1.8mm). For EEG recording, two stainless steel bones screws (MD-1310, BASi Research Products) were placed as EEG leads in the skull on the right frontal cortex (A/P 1mm, M/L -1mm), left parietal cortex (A/P -2mm, M/L -1mm). A third was placed on the cerebellum (A/P -6mm, M/L 0mm) for reference. Insulated wire leads were soldered to a headmount (8402, Pinnacle Technology) and wrapped around implanted screws for EEG recordings. Attached stainless steel wire leads were inserted directly into the neck musculature for EMG recordings. All cannula, screws, wires, and headmount were affixed to the skull using dental acrylic. Mice were placed into Raturn sampling cages (Bioanalytical Systems) in constant light.

### Biosensor Calibration

Prior to implantation, in vitro calibration of individual biosensors was performed in PBS at 37°, according to the manufacturer’s instructions and as previously described (25). Briefly, biosensors were attached to calibration test stage hardware (7052, Pinnacle Technology) and allowed to reach baseline. Three specific analyte injections (L-Glutamate, L-Lactate, and D-Glucose) and one interference injection (ascorbic acid) of a known concentration were introduced to the testing solution. All biosensors used in these experiments showed robust response to the specific analyte and no significant response to the interferents. The average response to each analyte injection was used as the calibration constant for that individual biosensor.

### Biosensor Recording

24 hours post-surgery, mice were briefly anesthetized for the implantation of two amperometric biosensors specific for either glucose, lactate, or glutamate (7004, Pinnacle Technology) into the guide cannula (n = 20 glucose, n = 12 lactate and glutamate). Biosensors were connected to a flexible preamplifier (100x amplification, EEG high pass filter: 0.5 Hz, EMG high pass filter: 10 Hz) and attached to the mouse’s headmount. The preamplifier was connected to a commutator, which passes electrical signal to the data acquisition system (8401, Pinnacle Technology) for sampling at 250 Hz for EEG/EMG channels and 1 Hz for biosensor channels. This set-up allowed the mouse unobstructed movement throughout the experiments. Recordings began after a stable baseline reading was reached using Sirenia Acquisition software. 24-hour recordings of uninterrupted sleep and biosensor data were used to establish a diurnal rhythm (n=6).

### Injection protocol and analysis

The following intraperitoneal (IP) injections were delivered to each mouse over a two day testing period: saline (0.9% NaCl, 2g/kg), glucose (50% dextrose, 2g/kg), and glibenclamide (Sigma-Aldrich, 1mg/kg dissolved in DMSO). Injections were time-locked across the light period (Day 1: 10am and 3pm, Day 2: 10am) and order of injections was randomized to account for any circadian rhythm influence. Biosensor data was exported from Sirenia software, combined into 10 second bins, and converted from nA to mM using the calibration constant for that biosensor. Data from the 10 minutes prior to the injection was used as a baseline measurement, while the 2 hours post-injection was analyzed relative to the established baseline. The average response to the saline controls was subtracted from the response to glucose or glibenclamide injections. Statistical analysis consisted of a one-way ANOVA with Dunnett’s post-hoc test to determine if the response was different from baseline and correlation analysis to assess the relationship between analytes.

### ISF Glucose and Lactate Power Spectra, Coherence, Phase Maps, and Cross Correlation Analysis

Individual biosensor power spectra, coherence, and phase maps were calculated via a multitaper approach, implemented using the *coherencyc* function from the open-source Chronux software package(29) with 95% confidence intervals (n=5). Power spectra were calculated across each glycemic state, binned into pre-defined frequencies, and compared using a two-way ANOVA with a correction for multiple comparisons. Cross correlations between glucose and lactate signals were calculated using cross-covariance function in MATLAB software. Briefly, the function compares z-scored values of lactate and glucose across varied time shifts. The significance of the relationship was evaluated using 95% confidence intervals.

### Sleep and Transition Analysis

EEG/EMG data were exported from the acquisition software and scored in 10-second epochs by hand according to conventional classifications for wake, NREM, and REM using Sirenia Sleep Software (n=10). Because sleep was only evaluated for 2 hours post-injection, there was not a significant amount of REM in most animals. Therefore, REM epochs were removed from all analyses. Change in sleep state was calculated by subtracting the percent time spent in each state following a saline injection from the individual glycemic challenges. To characterize transition data, sustained bouts (≥ 10 minutes for wake, ≥ 5 minutes for NREM, ≥1.5 minutes for REM) of each sleep state across a pre-defined peak response period were included (saline: 50-60 minutes post injection, glucose: 20-30 minutes post injection, glibenclamide: 35-45 minutes post injection, n=6). Analytes were converted to a percent change from the 30 seconds immediately preceding the transition. Statistical significance was evaluated using a one-way ANOVA with Dunnett’s post-hoc test.

### EEG Power Spectral Analyses

Power spectral analyses were calculated using Sirenia Sleep Pro Software (Pinnacle Technology) (n=10). Briefly, we used a Fast Fourier transform (Hann window) to calculate power spectra for all artifact-free epochs. Power spectral density was binned in 1 Hz increments and graphed by sleep state for each injection. For relative spectral power comparisons, data was separated into pre-defined frequency bins (Delta 0.5-4 Hz, Theta 4-8 Hz, Alpha 8-13 Hz, Sigma 12-15 Hz, Beta 13-30 Hz, Gamma 30-50 Hz, High Gamma 50-100 Hz) for each 10 second epoch. Relative power within each band was calculated as a percent of total power. Data was binned by sleep state to generate a state-dependent relative power spectrum for each mouse. Statistical differences were measured using 2-way ANOVA with post-hoc multiple comparisons and Tukey’s correction.

### EEG Variability and Coherence

EEG variability was calculated as root-mean square of each raw EEG signal over the same peak 10 minute bin as previously defined in transition analysis (n=10). Statistical differences were calculated using a one-way ANOVA with Dunnett’s post-hoc test. Coherence between the two raw EEG traces was calculated using Sirenia Sleep Pro software. Averages for each spectral band were calculated over the 10 minute peak response period. 2-way ANOVA with Tukey’s multiple comparisons correction for post-hoc analysis were used to assess differences across spectral bands.

### General Linear Modeling

We used general linear modeling (GLM) to evaluate the relationship between individual EEG spectral bands and ISF glucose or lactate levels as a function of sleep and glycemic state (n=6). The *glmfit* function in Matlab returns a beta coefficient for each condition according to a general linear model. One-sample t tests were used to determine the significance of each beta coefficient from zero.

### Lactate Injection Experiments

Surgeries and biosensor setup were completed as described for acute injection experiments. Mice (n=5) were given an injection of lactate (Sigma-Aldrich L1750, 50mg/kg dissolved in 0.9% saline) at the beginning of the light period. Biosensor and sleep data were analyzed two hours before injection for baseline and one hour after injection as previously described for acute injection experiments.

## Acknowledgements

We would like to acknowledge the following grants: F31AG066302 (CC), 1K01AG050719 (SLM), R01AG068330 (SLM), BrightFocus Foundation (A20201775S; SLM), Charleston Conference on Alzheimer’s disease New Vision Award (SLM), the McDonnell Center for Systems Neuroscience (SLM), P01NS080675 (DH, SLM, AM, MER), Averill Foundation (SLM).

## Data availability

All data generated during this study will be freely shared upon publication.

## Author Contributions

CMC, MS, SLM conceived of the study. CMC, MS, RVR, MER, DMH, RWG, KTK, SLM contributed to study design. CMC, MS, REI, AM, NJC, JAS, RVR, MER, DMH, RWG, KTK, SLM performed experiments, data analysis, and data interpretation. CMC, SLM wrote the manuscript. All authors discussed the results and commented on the manuscript.

**Supplementary Figure 1.**
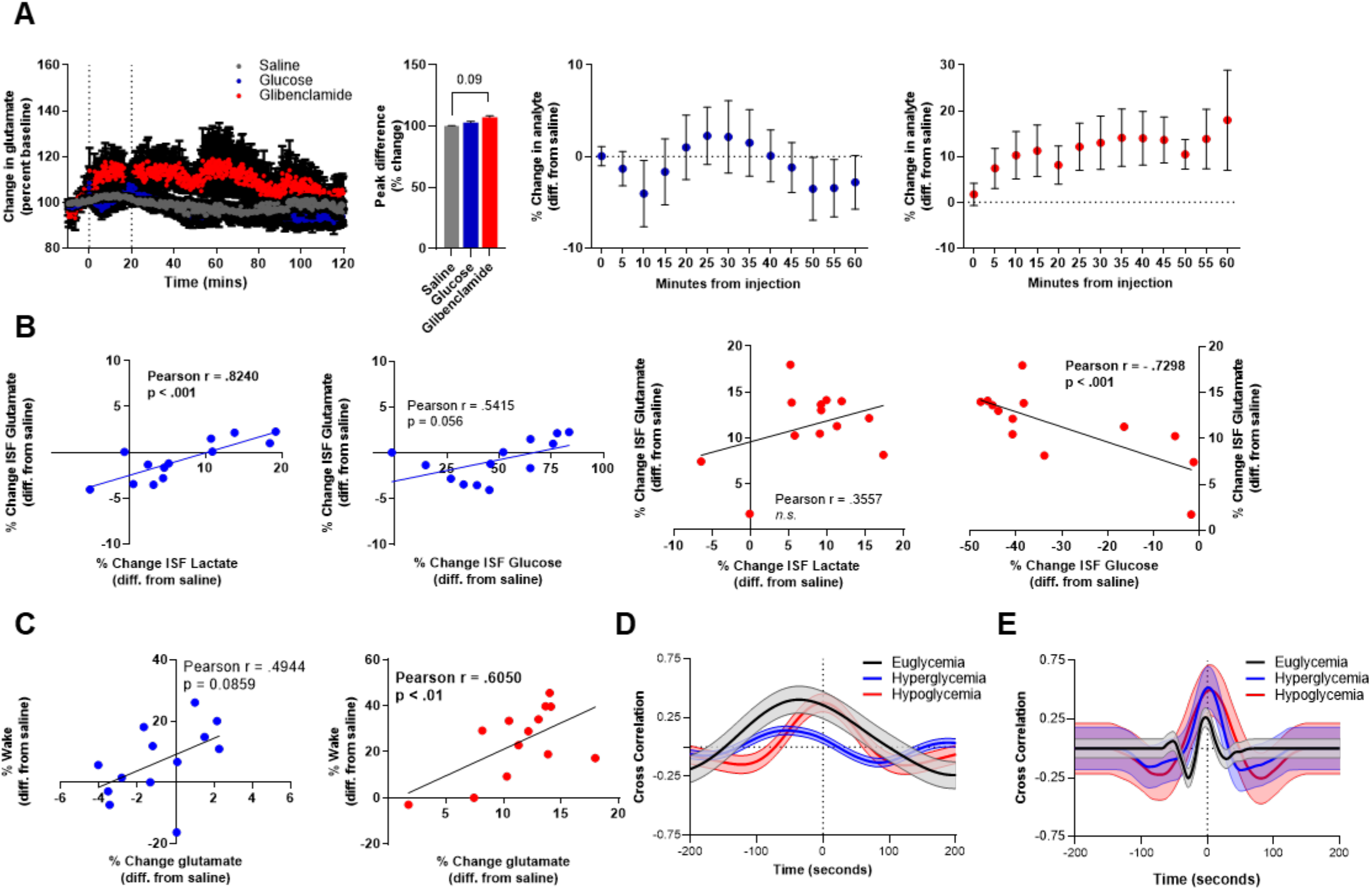
(**A**) ISF glutamate showed no differences from saline following glucose and glibenclamide injections. (**B**) Following glucose injections, increased ISF glutamate and lactate were positively correlated (*r =* .8240. p = 0.0005), while increased ISF glutamate and glucose trended towards a positive correlation (*r =* 0.5415. p = 0.056). During acute hypoglycemia, ISF glucose was negatively correlated with ISF glutamate (*r =* -0.7298. p = 0.0046). (**C**) During hyperglycemia, time spent awake trended towards a positive correlation with ISF glutamate *(r =* 0.4944. p = 0.0859), while during hypoglycemia, time awake was positively correlated with ISF glutamate (*r =* 0.6050. p < 0.01). (**D**) Cross correlation function shows a positive correlation between glucose and glutamate during all glycemic states. (**E**) The positive correlation between lactate and glutamate also persists across glycemic states. Data are reported as means ± SEM. *n* = 12 for injection. *n = 5 for x-correlation plots*. One-way ANOVA was used to evaluate significance between glycemic states.

**Supplementary Figure 2.**
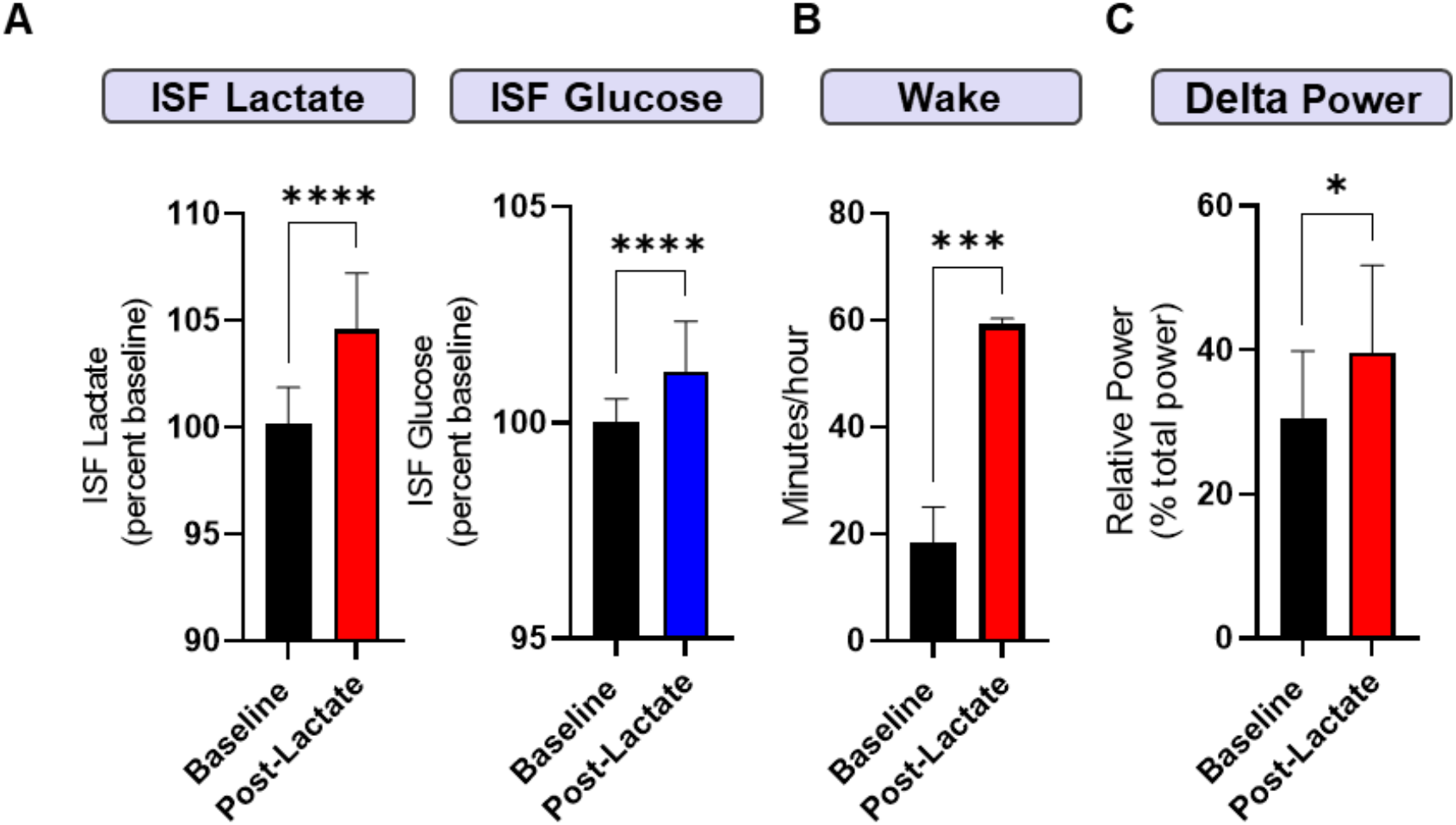
(**A**) Following an IP lactate injection, both ISF lactate (p<0.0001) and ISF glucose (p<0.0001) were elevated 15 minutes post injection. (**B**) Time spent awake was increased in the hour post lactate injection as compared to baseline. (**C**) Delta power increased following lactate injection, suggesting increased sleep propensity following this mild sleep deprivation. Data are reported as means ± SEM. *n* = *5. *p<0*.*05*, ^*#*^*p<0*.*01*, ^*^*^*p<0*.*001*, ^*+*^*p<0*.*0001*. Paired t tests were used to evaluate significance between baseline and post- injection.

